# Phylogenomics of the adaptive radiation of *Triturus* newts supports gradual ecological niche expansion towards an incrementally aquatic lifestyle

**DOI:** 10.1101/463752

**Authors:** B. Wielstra, E. McCartney-Melstad, J.W. Arntzen, R.K. Butlin, H.B. Shaffer

**Affiliations:** Department of Ecology and Evolutionary Biology, University of California, Los Angeles, CA 90095, USA.; Department of Animal and Plant Sciences, University of Sheffield, S10 2TN Sheffield, UK.; Naturalis Biodiversity Center, 2300 RA Leiden, The Netherlands.; Institute of Biology Leiden, Leiden University, 2300 RA, Leiden, The Netherlands.; La Kretz Center for California Conservation Science, Institute of the Environment and Sustainability, University of California, Los Angeles, CA 90095, USA.; Department of Marine Sciences, University of Gothenburg, Gothenburg 405 30, Sweden.

**Keywords:** morphology, phylogeny, sequence capture, systematics, target enrichment, transcriptome

## Abstract

Newts of the genus *Triturus* (marbled and crested newts) exhibit substantial variation in the number of trunk vertebrae (NTV) and a higher NTV corresponds to a longer annual aquatic period. Because the *Triturus* phylogeny has thwarted resolution to date, the evolutionary history of NTV, annual aquatic period, and their potential coevolution has remained unclear. To resolve the phylogeny of *Triturus*, we generated a c. 6,000 transcriptome-derived marker data set using a custom target enrichment probe set, and conducted phylogenetic analyses using: 1) data concatenation with RAxML, 2) gene-tree summary with ASTRAL, and 3) species-tree estimation with SNAPP. All analyses produce the same, highly supported topology, despite cladogenesis having occurred over a short timeframe, resulting in short internal branch lengths. Our new phylogenetic hypothesis is consistent with the minimal number of inferred changes in NTV count necessary to explain the diversity in NTV observed today. Although a causal relationship between NTV, body form, and aquatic ecology has yet to be experimentally established, our phylogeny indicates that these features have evolved together, and suggest that they may underlie the adaptive radiation that characterizes *Triturus*.

## 1. Introduction

Accurately retracing the evolution of phenotypic diversity in adaptive radiations requires well-established phylogenies. However, inferring the true branching order in adaptive radiations is hampered by the short time frame over which they typically unfold, which provides little opportunity between splitting events for phylogenetically informative substitutions to become established (resulting in low phylogenetic resolution; Philippe et al., 2011; Whitfield and Lockhart, 2007) and fixed (resulting in incomplete lineage sorting and discordance among gene-trees; Degnan and Rosenberg, 2006; Pamilo and Nei, 1988; Pollard et al., 2006). Resolving the phylogeny of rapidly multiplying lineages becomes even more complicated the further back in time the radiation occurred, because the accumulation of parallel substitutions along terminal branches can lead to long-branch attraction (Felsenstein, 1978; Swofford et al., 2001). A final impediment is reticulation between closely related (and not necessarily sister-) species through past or ongoing hybridization, resulting in additional gene-tree/species-tree discordance (Kutschera et al., 2014; Leaché et al., 2014; Mallet et al., 2016).

Phylogenomics, involving the consultation of a large number of markers spread throughout the genome, has proven successful in resolving both recent (e.g. Giarla and Esselstyn, 2015; Leaché et al., 2016; Léveillé-Bourret et al., 2018; Meiklejohn et al., 2016; Nater et al., 2015; Scott et al., 2018; Shi and Yang, 2018) and more ancient (e.g. Crawford et al., 2012; Irisarri and Meyer, 2016; Jarvis et al., 2014; McCormack et al., 2012; Song et al., 2012) evolutionary radiations. The appeal of greatly increasing the amount of data available for any given phylogenetic problem is that it often (but not always; see Philippe et al., 2011) provides informative characters to resolve short branches in the tree of life. Advances in laboratory and sequencing techniques, bioinformatics, and tree-building methods all facilitate phylogenetic reconstruction based on thousands of homologous loci for a large number of individuals, and promise to help provide the phylogenetic trees necessary to interpret the evolution of eco-morphological characters involved in adaptive radiations (Alföldi et al., 2011; Stroud and Losos, 2016). In this study, we conduct a phylogenomic analysis of an adaptive radiation that moderately-sized multilocus nuclear DNA datasets (Arntzen et al., 2007; Espregueira Themudo et al., 2009; Wielstra et al., 2014) have consistently failed to resolve: the Eurasian newt genus *Triturus* (Amphibia: Urodela: Salamandridae), commonly known as the marbled and crested newts.

One of the most intriguing features of *Triturus* evolution is the correlation between certain aspects of their ecology and the number of trunk vertebrae (NTV; Fig. 1). Species characterized by a higher modal NTV (which translates into a more elongate body build with proportionally shorter limbs) are associated with a more aquatic lifestyle. Empirically, the number of months a *Triturus* species spends in the water (defined at the population level as the peak date of emigration, leaving a breeding pond, minus the peak in immigration, entering it) roughly equals NTV minus 10 (Arntzen, 2003; Arntzen and Wallis, 1999; Slijepčević et al., 2015). The intrageneric variation in NTV shown by *Triturus*, ranging from 12 to 17, is unparalleled in the family Salamandridae (Arntzen et al., 2015; Lanza et al., 2010) and a causal relationship between NTV expansion and an increasingly aquatic lifestyle has been presumed, but never adequately placed into a phylogenetic comparative analysis (Arntzen, 2003; Arntzen et al., 2015; Arntzen and Wallis, 1999; Govedarica et al., 2017; Slijepčević et al., 2015; Urošević et al., 2016; Vukov et al., 2011; Wielstra and Arntzen, 2011). A well-established *Triturus* species-tree is required to accurately retrace NTV evolution and assess the concordance between aquatic lifestyle and NTV across the genus.

**Fig. 1.**
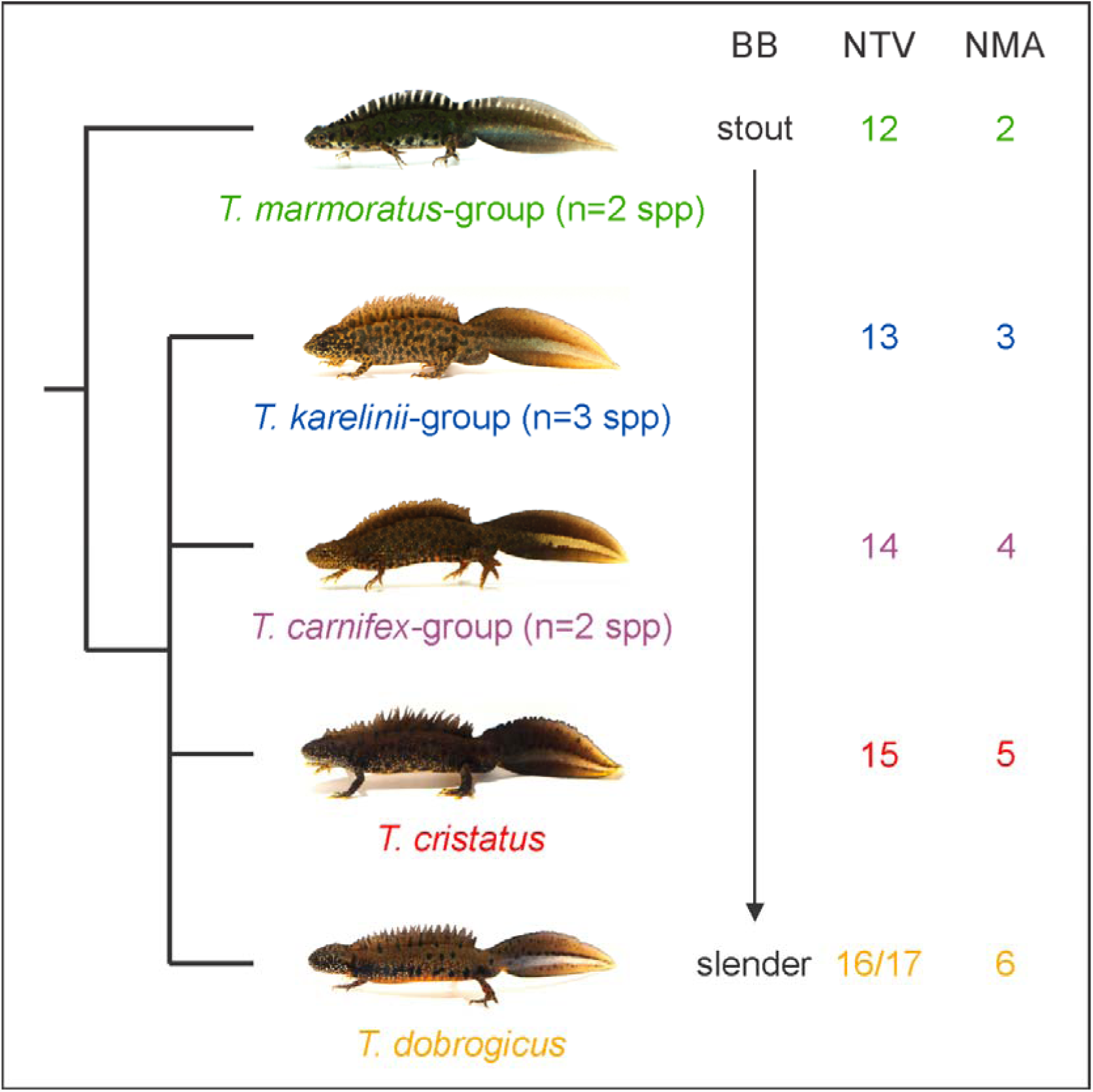
The adaptive radiation of *Triturus* newts. Five body builds (BB) from stout to slender are observed in *Triturus* that are also characterized by an increasing number of trunk vertebrae (NTV) and number of annual aquatic months (NMA). The marbled newts (*T*. *marmoratus*-group) and crested newts (remaining four BBs) are sister clades. Relationships among the crested newts are not yet resolved and are the main focus of the present study.

Our goal is to obtain a genome-enabled phylogeny for *Triturus* and use it to reconstruct the eco-morphological evolution of NTV and aquatic/terrestrial ecology across the genus. As the large size of salamander genomes hampers whole-genome sequencing (but see Elewa et al., 2017; Nowoshilow et al., 2018; Smith et al., 2018), we employ a genome-reduction approach in which we capture and sequence a set of transcriptome-derived markers using target enrichment, an efficient technique that affords extremely high resolution at multiple taxonomic levels (Abdelkrim et al., 2018; Bi et al., 2012; Bragg et al., 2016; Gnirke et al., 2009; McCartney-Melstad et al., 2016; McCartney-Melstad et al., 2018). Using data concatenation (with RAxML), gene-tree summarization (with ASTRAL) and species-tree estimation (with SNAPP), we fully resolve the *Triturus* phylogeny and place the extreme body shape and ecological variation observed in this adaptive radiation into an evolutionary context.

## 2. Materials and Methods

### 2.1. Target capture array design

Nine *Triturus* newts (seven crested and two marbled newt species) and one banded newt (*Ommatotriton*) were subjected to transcriptome sequencing. Transcriptome assemblies for each species were generated using Trinity v2.2.0 (Grabherr et al., 2011), clustered at 90% using usearch v9.1.13 (Edgar, 2010), and subjected to reciprocal best blast hit analysis (Bork et al., 1998; Camacho et al., 2009; Tatusov et al., 1997) to produce a set of *T*. *dobrogicus* transcripts (the species with the highest quality transcriptome assembly) that had putative orthologues present in the nine other transcriptome assemblies. These transcripts were then annotated using blastx to *Xenopus tropicalis* proteins, retaining one annotated transcript per protein. We attempted to discern splice sites in the transcripts, as probes spanning splice boundaries may perform poorly (Neves et al., 2013), by mapping transcripts iteratively to the genomes of *Chrysemys picta* (Shaffer et al., 2013), *X*. *tropicalis* (Hellsten et al., 2010), *Nanorana parkerii* (Sun et al., 2015) and *Rana catesbeiana* (Hammond et al., 2017). A single exon ≥ 200bp and ≤ 450bp was retained for each transcript target. To increase the ability of the target set to capture markers across all *Triturus* species, orthologous sequences from multiple species were included for targets with > 5% sequence divergence from *T*. *dobrogicus* (Bi et al., 2012). We generated a target set of 7,102 genomic regions for a total target length of approximately 2.3 million bp. A total of 39,143 unique RNA probes were synthesized as a MyBaits-II kit for this target set at approximately 2.6X tiling density by Arbor Biosciences (Ann Arbor, MI, Ref# 170210-32). A detailed outline of the target capture array design process is presented in Supplementary Text S1.

### 2.2. Sampling scheme

We sampled 23 individual *Triturus* newts (Fig. 2; Supplementary Table S1) for which tissues were available from previous studies (Wielstra et al., 2017a; Wielstra et al., 2017b; Wielstra et al., 2013). Because the sister-group relationship between the two marbled and seven crested newts is well established (Fig. 1), while the relationships among the crested newt species have defied resolution, we sampled the crested newt species more densely, including three individuals per species to include intraspecific differentiation and to avoid misleading phylogenies resulting from single exemplar sampling (Spinks et al., 2013). Because *Triturus* species show introgressive hybridization at contact zones (Arntzen et al., 2014), we aimed to reduce the impact of interspecific gene flow by only including individuals that originate away from hybrid zones and have previously been interpreted as unaffected by interspecific genetic admixture (Wielstra et al., 2017a; Wielstra et al., 2017b). The reality of phylogenetic distortion by interspecific gene flow was underscored in a test for the phylogenetic utility of the transcripts used for marker design which included a genetically admixed individual (details in Supplementary Text S1).

**Fig. 2.**
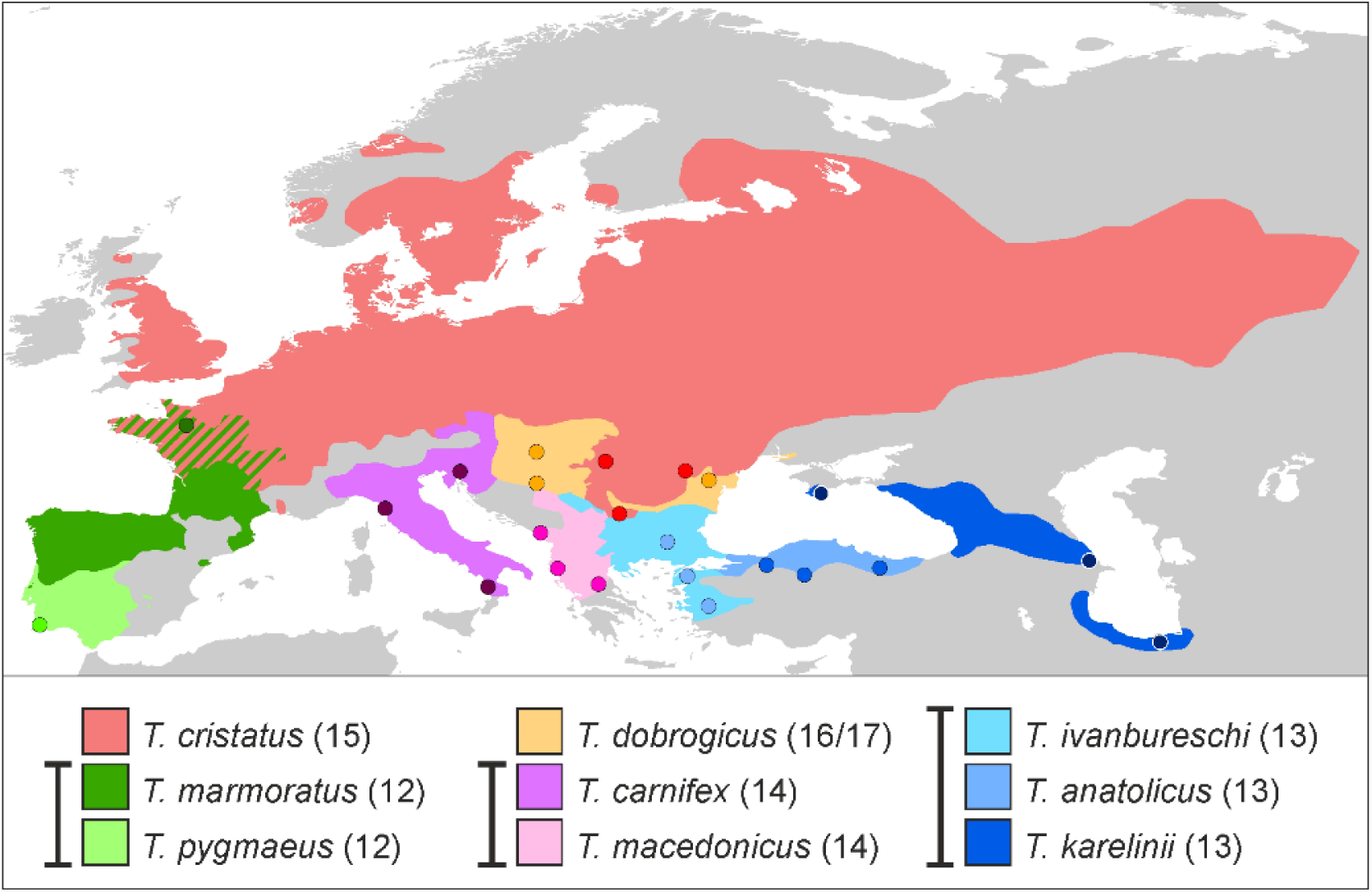
Distribution and sampling scheme for *Triturus*. Dots represent sample localities (details in Supplementary Table S1). For the marbled newts (in green) a single individual is sampled for each of the two species and for the crested newts (other colours) three individuals are sampled for all seven species. The number in parentheses reflects each species’ characteristic number of trunk vertebrae and whiskers link species that possess the same body build (see Fig. 1).

### 2.3. Laboratory methods

DNA was extracted from samples using a salt extraction protocol (Sambrook and Russell, 2001), and 10,000ng per sample was sheared to approximately 200bp-500bp on a BioRuptor NGS (Diagenode) and dual-end size selected (0.8X-1.0X) with SPRI beads. Dual-indexed libraries were prepared from 375-2000ng of size selected DNA using KAPA LTP library prep kits (Glenn et al., 2017). These libraries were pooled (with samples from other projects) into batches of 16 samples at 250ng per sample (4,000ng total) and enriched in the presence of 30,000ng of c0t-1 repetitive sequence blocker (McCartney-Melstad et al., 2016) derived from *T*. *carnifex* (casualties from a removal action of an invasive population (Meilink et al., 2015)) by hybridizing blockers with libraries for 30 minutes and probes with libraries/blockers for 30 hours. Enriched libraries were subjected to 14 cycles of PCR with KAPA HiFi HotStart ReadyMix and pooled at an equimolar ratio for 150bp paired-end sequencing across multiple Illumina HiSeq 4000 lanes (receiving an aggregate of 18% of one lane, for a multiplexing equivalent of 128 samples per lane).

### 2.4. Processing of target capture data

A total of 3,937,346 read pairs from the sample receiving the greatest number of reads were used to *de novo* assemble target sequences for each target region using the assembly by reduced complexity (ARC) pipeline (Hunter et al., 2015). A single assembled contig was selected for each original target region by means of reciprocal best blast hit (RBBH) (Rivera et al., 1998), and these were used as a reference assembly for all downstream analyses. Adapter contamination was removed from sample reads using skewer v0.2.2 (Jiang et al., 2014), and reads were then mapped to the reference assembly using BWA-MEM v0.7.15-r1140 (Li, 2013). Picard tools v2.9.2 (https://broadinstitute.github.io/picard/) was used to add read group information and to mark PCR duplicates, and HaplotypeCaller and GenotypeGVCFs from GATK v3.8 (McKenna et al., 2010) were used jointly to genotype the relevant groups of samples (either crested newts or crested newts + marbled newts depending on the analysis; see below). SNPs that failed any of the following hard filters were removed: QD < 2, MQ < 40, FS > 60, MQRankSum < −12.5, ReadPosRankSum < −8, and QUAL < 30 (Poplin et al., 2017). We next attempted to remove paralogous targets from our dataset with a Hardy Weinberg Equilibrium (HWE) filter for heterozygote excess. Heterozygote excess p-values were calculated for every SNP using vcftools 0.1.15 (Danecek et al., 2011), and any target containing at least one SNP with a heterozygote excess p-value < 0.05 was removed from downstream analysis. More detail on the processing of the target capture data can be found in Supplementary Text S2.

### 2.5. Phylogenetic analyses

A concatenated maximum likelihood phylogeny was inferred with RAxML version 8.2.11 (Stamatakis, 2014) based on an alignment of 133,601 SNPs across 5,866 different targets. We included all 23 *Triturus* individuals in this analysis. For gene-tree summary, ASTRAL v5.6.1 (Zhang et al., 2017) was used to estimate the crested newt species-tree from 5,610 gene-trees generated in RAxML. The 21 crested newt samples were assigned species membership, and no marbled newts were included because estimating terminal branch lengths is not possible for species with a single representative. For species-tree estimation, SNAPP v1.3.0 (Bryant et al., 2012) within the BEAST v2.4.8 (Bouckaert et al., 2014) environment was used to infer the crested newt species-tree from single biallelic SNPs randomly selected from each of 5,581 post-filtering targets. All three individuals per crested newt species were treated as a single terminal and marbled newts were again excluded given our single exemplar sampling of both species. We also estimated divergence times in SNAPP for the crested newts. The split between *T*. *carnifex* and *T*. *macedonicus*, assumed to correspond to the origin of the Adriatic Sea at the end of the Messinian Salinity Crisis 5.33 million years ago, was used as a single calibration point (Arntzen et al., 2007; Wielstra and Arntzen, 2011) to produce a rough estimate of the timing of cladogenesis. A detailed description of our strategy for phylogenetic analyses is available in Supplementary Text S3.

## 3. Results

Samples received a mean of 2,812,980 read pairs (s.d. = 585,815). Enrichment was highly efficient, especially given the large genome size of *Triturus*, with an average of 44.5% of raw reads mapping to the assembled target sequences (s.d. = 2.6%). After removing PCR duplicates, which accounted for an average of 22.6% of mapped reads, the unique read on target rate was 34.4% (s.d. = 1.9%). The 23 samples in the final RAxML alignment contained an average of 10.1% missing data (min = 3.2%, max = 31.8%) after setting genotype calls with GQ scores of less than 20 to missing.

The concatenated analysis with RAxML supports a basal bifurcation in *Triturus* between the marbled and crested newts (Fig. 3), consistent with the prevailing view that they are reciprocally monophyletic (Arntzen et al., 2007; Espregueira Themudo et al., 2009; Wielstra et al., 2014). RAxML also recovers each of the crested newt species as monophyletic, validating our decision to collapse the three individuals sampled per species in a single terminal in ASTRAL and SNAPP. Furthermore, all five *Triturus* body builds are recovered as monophyletic (cf. Arntzen et al., 2007; Espregueira Themudo et al., 2009; Wielstra et al., 2014). The greatest intraspecific divergence is observed in *T*. *carnifex* (Supplementary Text S1; Supplementary Fig. S1; Supplementary Table S2).

**Fig. 3.**
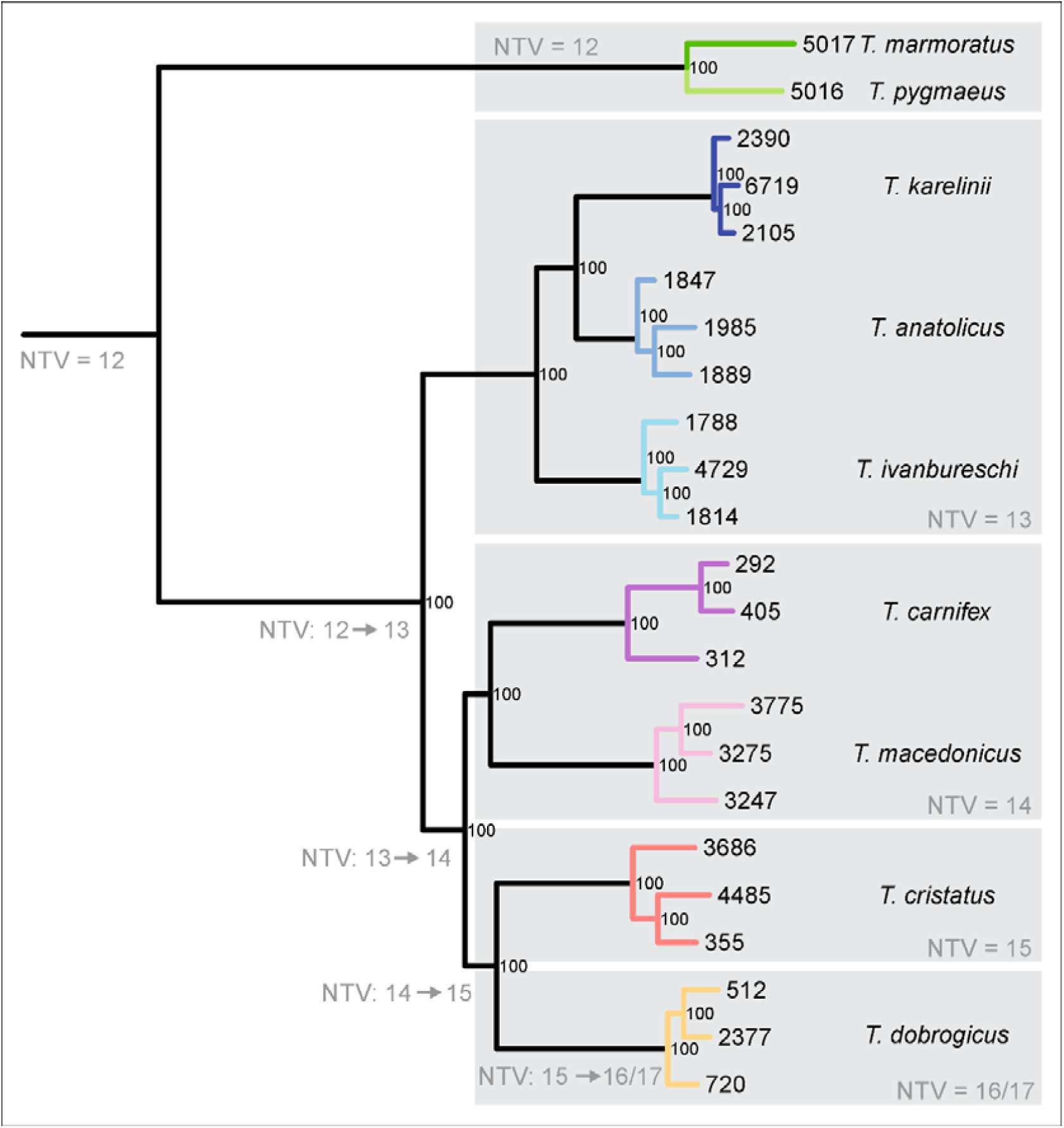
*Triturus* newt phylogeny based on data concatenation with RAxML. This maximum likelihood phylogeny is based on 133,601 SNPs derived from 5,866 nuclear markers. Numbers at nodes indicate bootstrap support from 100 rapid bootstrap replicates. The five *Triturus* body builds (see Fig. 1) are delineated by grey boxes, with their characteristic number of trunk vertebrae (NTV) noted. Inferred changes in NTV under the parsimony criterion are noted along branches. Colours reflect species and correspond to Fig. 2. Tip labels correspond to Supplementary Table S1.

Phylogenetic inference based on data concatenation with RAxML (Fig. 3), gene-tree summary with ASTRAL (Fig. 4a) and species-tree estimation with SNAPP (Fig. 4b) all recover the same crested newt topology, with a basal bifurcation between the *T*. *karelinii*-group (NTV = 13; *T*. *ivanbureschi* as the sister taxon to *T*. *anatolicus* + *T*. *karelinii)* and the remaining taxa, which themselves are resolved into the species pairs *T*. *carnifex* + *T*. *macedonicus* (NTV=14; the *T*. *carnifex*-group), and *T*. *cristatus* (NTV=15) + *T*. *dobrogicus* (NTV=16/17). Despite the rapidity of cladogenesis, we obtain strong branch support for every internal node. Even with the uncertainty in dating given a single biogeographically-derived calibration date, the bifurcation giving rise to the four crested newt species groups (cf. Fig. 1) must have occurred over a relatively short time frame (Fig. 5), reflected by two particularly short, but resolvable internal branches (Fig. 3; Fig. 4).

**Fig. 4.**
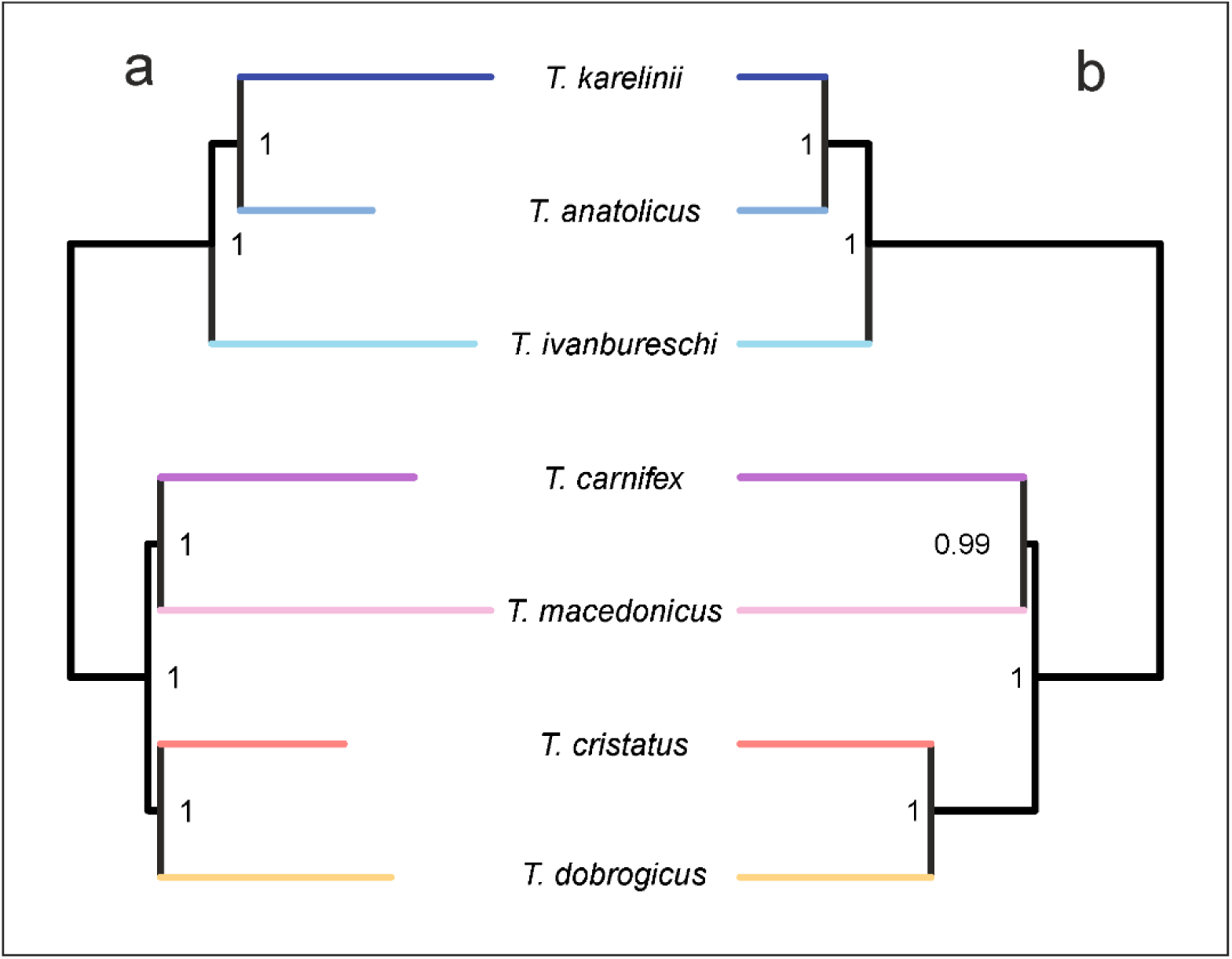
Crested newt phylogeny based on gene-tree summary with ASTRAL and species-tree estimation with SNAPP. The ASTRAL tree (a) is based on 5,610 gene-trees. Numbers at nodes indicate local quartet support posterior probabilities. The SNAPP tree (b) is based on single biallelic SNPs taken from 5,581 nuclear markers. Numbers at nodes indicate posterior probabilities. Colours reflect species and correspond to Fig. 2. Note that both topologies are identical to the phylogeny based on data concatenation (Fig. 3).

**Fig. 5.**
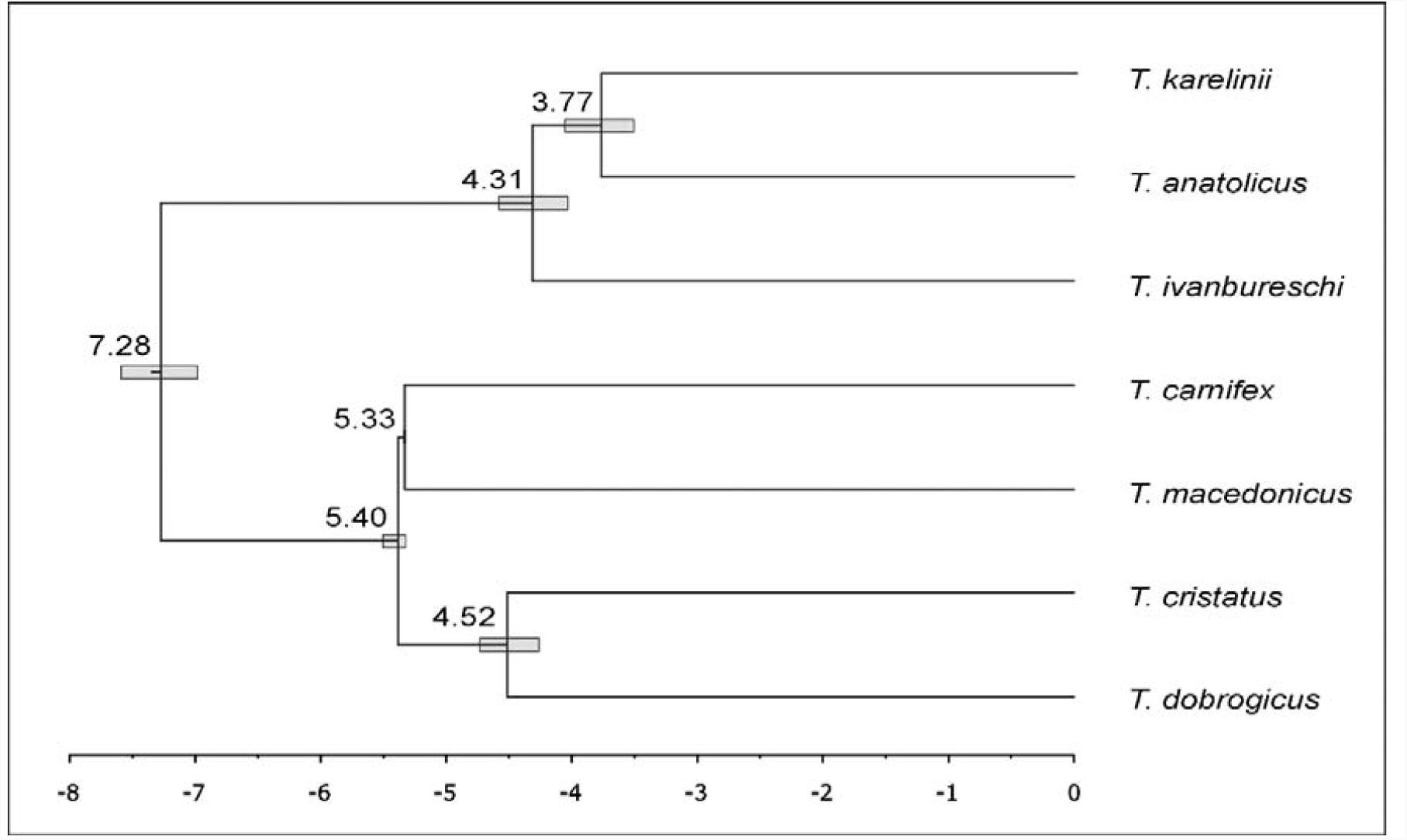
Dated species-tree for the crested newts. Divergence times were determined with SNAPP, using a single *T*. *carnifex*–*T*. *macedonicus* inferred split date of 5.33 million years ago as a calibration point. Numbers at nodes reflect median divergence times in millions of years ago and bars the 95% credibility interval around the median.

The phylogenomic analyses suggest considerable gene-tree/species-tree discordance in *Triturus*. The normalized quartet score of the ASTRAL tree (Fig. 4a), which reflects the proportion of input gene-tree quartets consistent with the species-tree, is 0.63, indicating a high degree of gene-tree discordance. Furthermore, the only node in the SNAPP tree with a posterior probability below 1.0 (i.e. 0.99) is subtended by a very short branch (Fig. 4b). Consistent with the high level of gene-tree/species-tree discordance, we also found that the full mtDNA-based phylogeny of *Triturus* produced a highly supported, but topologically different, phylogeny (Supplementary Text S3; Supplementary Fig. S2; Wielstra and Arntzen, 2011).

Considering an NTV count of 12, as observed in the marbled newts as well as the most closely related newt genera, as the ancestral state for *Triturus* (Arntzen et al., 2015; Veith et al., 2018), three sequential single-vertebral additions to NTV along internal branches, and one or two additions along the terminal branch leading to *T*. *dobrogicus* (in which NTV = 16 and NTV = 17 occur at approximately equal frequency; Arntzen et al., 2015; Wielstra et al., 2016) are required under a parsimony criterion (with either ACCTRAN or DELTRAN optimization) to explain the present-day variation in NTV observed in *Triturus* (Fig. 3). This is the minimum possible number of inferred changes in NTV count required to explain the NTV radiation observed today (Supplementary Fig. S3; Supplementary Text S5). No NTV deletions or reversals are required, implying a linear, stepwise, single-addition scenario for NTV expansion in *Triturus*.

## 4. Discussion

We use a large, tramscriptome-derived phylogenomic dataset to construct a phylogenetic hypothesis and study the evolution of ecological and phenotypic diversity within the adaptive radiation of *Triturus* newts. In contrast to previous attempts to recover a multilocus species-tree (Arntzen et al., 2007; Espregueira Themudo et al., 2009; Wielstra et al., 2014), we recover full phylogenetic resolution with strong support across the tree. Despite cladogenesis having occurred in a relatively brief time window (Fig. 5), resulting in a high degree of gene-tree/species-tree discordance, independent phylogenetic approaches based on data concatenation (RAxML), gene-tree summarization (ASTRAL) and species-tree estimation (SNAPP), all recover the same, highly supported topology for *Triturus* (Fig. 3; Fig. 4). Our *Triturus* case study underscores that sequence capture by target enrichment is a promising approach to resolve the phylogenetic challenges associated with adaptive radiations, particularly for taxa with large and complicated genomes where other genomic approaches are impractical, including salamanders (McCartney-Melstad et al., 2016).

Our new phylogenetic hypothesis allows us to place the eco-morphological differentiation shown by *Triturus* into a coherent evolutionary context. Over time, *Triturus* expanded its range of NTV to encompass higher counts (Fig. 3). The *Triturus* tree is consistent with a maximally parsimonious scenario, under which four to five character state changes are required to explain the radiation in NTV observed today. Any other possible phylogenetic relationship among *Triturus* body builds would require a higher number of inferred NTV changes (Supplementary Fig. S3). Three of these inferred changes are positioned along internal branches, of which two are particularly short, suggesting that changes in NTV count can evolve over a relatively short time. The fourth and fifth inferred change are situated on the external branch leading to *T*. *dobrogicus*, the only *Triturus* species with substantial intraspecific variation in NTV count (Arntzen et al., 2015; Wielstra et al., 2016).

Newts annually alternate between an aquatic and a terrestrial habitat, and the functional trade-off between adaptation to life in water or on land likely poses contrasting demands on body build (Fish and Baudinette, 1999; Gillis and Blob, 2001; Gvo□dík and van Damme, 2006; Shine and Shetty, 2001). Considering the observed relationship between one additional trunk vertebra and an extra month annually spent in the water (Fig. 1), the extraordinary NTV variation observed in *Triturus* may reflect the morphological mechanism by which more efficient exploitation of a wider range in hydroperiod (i.e. the annual availability of standing water) evolved. Despite the evolvability of NTV count (Arntzen et al., 2015), NTV evolution has been phylogenetically constrained in *Triturus*. Apparently the change in NTV was directional and involved the addition of a single trunk vertebra at a time (Fig. 3; Supplementary Fig. S3). Species with a more derived body build, reflected in a higher NTV, have a relatively prolonged aquatic period and, because species with transitional NTV counts remain extant, the end result is an eco-morphological radiation.

*Triturus* newts show a slight degree of intraspecific variation in NTV today. Such variation is partially explained by interspecific hybridization (emphasizing the genetic basis of NTV count; Arntzen et al., 2014), but there is standing variation in NTV count within all *Triturus* species (Slijepčević et al., 2015). This suggests that, during *Triturus* evolution, there has always been intraspecific NTV count polymorphism that could be subjected to natural selection. Whether there is a causal relationship between the directional, parsimonious evolution of higher NTV and the equally parsimonious evolutionary increase in aquatic lifestyle, and, if so, which of these two may be the actual target of selection, remain important open questions. A proper understanding of the functional relationship between NTV, body build and fitness in aquatic/terrestrial environments in *Triturus* is still lacking (Gvo□dík and van Damme, 2006), and functional studies exploring this fitness landscape across intra and interspecific variation in NTV is an important next step in establishing a firm causal relationship between variation, performance and fitness. The recent availability of the first salamander genomes (Elewa et al., 2017; Nowoshilow et al., 2018; Smith et al., 2018) finally offers the prospect of sequencing the genome of each *Triturus* species and exploring the developmental basis for NTV and its functional consequences in the diversification of the genus.

## Acknowledgements

Andrea Chiocchio, Daniele Canestrelli, Michael Fahrbach, Ana Ivanović, Raymond van der Lans, and Kurtuluş Olgun helped obtain samples for transcriptome sequencing. Permits were provided by the Italian Ministry of the Environment (DPN-2009-0026530), the Environment Protection Agency of Montenegro (no. UPI-328/4), the Ministry of Energy, Development and Environmental Protection of Republic of Serbia (no. 353-01-75/2014-08), and TÜBİTAK, Turkey (no. 113Z752). RAVON & Natuurbalans-Limes Divergens provided the *T*. *carnifex* used to create c0t-1. Tara Luckau helped in the lab. Peter Scott provided valuable suggestions on methodology. Wiesław Babik and Ana Ivanović commented on an earlier version of this manuscript. This work used the Vincent J. Coates Genomics Sequencing Laboratory at UC Berkeley, supported by NIH S10 Instrumentation Grants S10RR029668 and S10RR027303. Computing resources were provided by XSEDE (Towns et al., 2014) and the Texas Advanced Computing Center (TACC) Stampede2 cluster at The University of Texas at Austin.

## Data availability

Raw sequence read data for the sequence capture libraries of the 23 *Triturus* samples and the 12 transcriptome libraries are available at SRA (PRJNA498336). Transcriptome assemblies, genotype calls (VCF) for the 21- and 23-sample datasets, input files for the RAxML, ASTRAL and SNAPP analyses, and synthesized target sequences are available at Zenodo (https://doi.org/10.5281/zenodo.1470914). Supplementary data associated with this article can be found, in the online version, at [xxx]

