## Supplemental text, figures, tables for "Phylogenomics of the adaptive radiation of *Triturus* newts supports gradual ecological niche expansion towards an incrementally aquatic lifestyle"

**Supplementary Text, Figures and Tables**

**Supplemental Text S1-S3**

**Text S1: Array Design**

*Transcriptome sequencing* – Liver tissue samples in RNAlater from ten newts (one each of *Triturus anatolicus*, *T. carnifex*, *T. cristatus*, *T. dobrogicus*, *T. ivanbureschi*, *T. karelinii*, *T. macedonicus*, *T. marmoratus*, *T. pygmaeus*, and *Ommatotriton nesterovi*; Supplementary Table S1) were sent to ZF-Genomics (Leiden, The Netherlands) for RNA extraction and sequencing on a HiSeq 2500. Samples received an average of 43,810,415 clusters (SD=9,744,176) in 150bp paired-end configuration.

*QC and Assembly* – Paired-end sequencing reads were trimmed for adapter contamination and sequence quality using a 4-bp sliding window in Trimmomatic 0.32 (Bolger, et al. 2014), clipping the 3’ ends of reads when the average sequence quality within the window dropped below 20. Leading bases with a quality score less than 5 and trailing bases with a quality score less than 15 were also removed, and reads shorter than 40bp after trimming were discarded.

A median of 38,575,204 read pairs were input into the Trinity assembler for each of the ten species (min=27,572,854, max=54,993,188, sd=8,916,227), and a median of 18.6% of these were retained after *in silico* normalization (min=15.8%, max=22.8%, sd=2.3%). Each transcriptome was individually assembled using Trinity 2.2.0 with read coverage normalized to a maximum of 50 (Grabherr, et al. 2011). Individual Trinity assemblies were clustered at 90% identity using usearch v9.1.13 to reduce redundancy (Edgar 2010). Assemblies contained a median of 157,608 contigs after clustering at 90% similarity (min=80,803 for *T. karelinii*, and max=182,488 for *T. carnifex*).

These clustered assemblies were then used for pairwise comparison between *T. dobrogicus* and the other nine species using *blastn* v2.2.30 (Camacho, et al. 2009). The reciprocal best blast hits (RBBH) method was used to determine presumptive orthology between the assembled transcripts for each pairwise species comparison (Tatusov, et al. 1997; Bork, et al. 1998). *T. dobrogicus* transcripts that returned reciprocal best blast hits to all of the nine other species were retained and all other transcripts were discarded.

*Transcriptome comparison* – The remaining set of 10,333 *T. dobrogicus* transcripts was self-blasted to attempt to reduce redundancy, which may help reduce the inclusion of multiple isoforms of the same gene, chimeric transcripts assembled by Trinity, and transcripts with truly similar regions that may complicate downstream bioinformatics. As a conservative measure, both the subject and query transcript were discarded if any transcript showed significant similarity (blast e-value < 0.001) to a different transcript or to different regions of itself.

*Annotation* – The remaining set of 9,214 *T. dobrogicus* transcripts were annotated using a translated blastx search to known *X. tropicalis* proteins with an e-value cutoff of 0.1 (Hellsten, et al. 2010). Transcripts that did not have a positive blastx hit to the *Xenopus* protein database were discarded, and only a single transcript annotating to a particular *Xenopus* protein was retained.

*Splice site prediction* – For the remaining set of 7,228 *T. dobrogicus* transcripts we attempted to infer splice sites in the candidate targets to avoid designing baits that span such boundaries, as these baits may perform poorly (Neves, et al. 2013) and because targeting a single exon for each transcript simplifies downstream analyses. Splice sites were predicted by attempting to map each transcript to the *Chrysemys picta* genome (Shaffer, et al. 2013) using exonerate’s est2genome model (Slater and Birney 2005) with a DNA word length of 10. Approximately 93% of all transcripts (n=6,758) successfully mapped to the *C. picta* genome, and for regions that mapped, the longest contiguous section of the mapped transcript was harvested. If the longest contiguous segment was less than 200bp, the first high-scoring segment pair (HSP) was extended towards the 5’ end until reaching 200bp, followed by extending the final HSP towards the 3’ end until reaching 200bp if necessary. Of the 6,758 transcripts that mapped to *C. picta*, 69 transcripts did not have an HSP longer than 200bp and could not be extended to 200bp in the 5’ or 3’ direction and were dropped as prospective targets.

The 470 transcripts that did not align to the *C. picta* genome were sequentially aligned to the genomes of *X. tropicalis* (Hellsten, et al. 2010), *Nanorana parkerii* (Sun, et al. 2015), and *Rana catesbeiana* (Hammond, et al. 2017) to attempt to find splice sites, taking the first successful species alignment from the list. Of these 470 transcripts, 125 mapped to *X. tropicalis*, 39 mapped to *N. parkerii*, and 36 mapped to *R. catesbeiana*. Again the longest contiguous aligned segment of each transcript was retained as a possible target, and transcripts with no aligned HSP of at least 200bp had their alignments extended in the 5’ then 3’ directions to attain targets of at least 200bp. For the 270 transcripts that did not map to any genome, the first (leftmost) 300bp of the assembled transcript was selected for a target region (except for the one transcript that was only 231bp long—for this target the entire 231bp transcript was used). It is possible that this leftmost orientation may enrich these targets for UTR sequence, assuming that the transcript was fully assembled by Trinity.

All exon targets were trimmed to a maximum of 450bp (from the 3’ edge) and checked again for complementarity using a self BLAST in blastn. The first qualifying target from each unique Trinity cluster-gene identifier was retained, and any other targets that arose from the same Trinity gene identifier were discarded (n=19). This target set contains sub-sequences from 7,139 different transcripts for a total length of 2,272,851bp (mean of each sub-sequence=318bp, min=200bp, max=400bp, median=300bp).

As we are interested in capturing these loci from all *Triturus* taxa, including both crested and marbled newts, we decided to include probes designed from multiple species for the same target if divergence between representative species in the two main clades was greater than 5% (Bi, et al. 2012). Since the bulk of the target sequences were designed from *T. dobrogicus*, which together with *T. carnifex*, *T. cristatus* and *T. macedonicus* encompasses one of two main clades in the crested newts (Wielstra and Arntzen 2011; Wielstra, Arntzen, et al. 2014), the three remaining species of crested newts encompassing the other clade (*T. karelinii*, *T. anatolicus, and T. ivanbureschi*) were used to determine if greater than 5% divergence existed between the two major clades for that target. First, the *T. dobrogicus* targets were blasted against *T. karelinii*, enforcing a full-length HSP with respect to the query sequence, yielding 2,850 hits; 30 of these were found to have a divergence greater than 5% and were added to the 7,139 *T. dobrogicus* targets. Then the remaining 4,289 *T. dobrogicus* targets were blasted to *T. anatolicus*, yielding 2,883 hits and an additional 35 targets. Finally the remaining 1,406 *T. dobrogicus* targets were blasted to *T. ivanbureschi*, yielding 631 hits and 10 more targets. Subsequently the process was repeated for the marbled newts *T. pygmaeus* and *T. marmoratus*, which constitute the sister lineage of the two crested newt clades, yielding an additional 222 and 27 targets after positive hits for 5,544 of 7,139 targets and 440 of 1,595 residual targets, respectively. Overall, an additional 324 orthologous targets that were more than 5% divergent between *T. dobrogicus* and other *Triturus* species were added to attempt to generate a set of probes that would perform well across the genus.

A set of 7,463 target sequences (average length=317bp, min=175bp, max=474bp) was sent to Arbor Biosciences for probe tiling and synthesis. After removing any probes softmasked by RepeatMasker and the Amphibia database, 39,143 unique 120 bp RNA probes were synthesized at approximately 2.6X tiling density across 7,418 target sequences by Arbor Biosciences (Ann Arbor, MI) as a MyBaits-II kit.

*Test for phylogenetic utility* – The phylogenetic utility of the genomic transcript markers was validated by building a phylogeny from the transcript sequences with RAxML. Trinity-assembled transcriptomes were clustered at 90% identity using usearch v9.1.13 (Edgar 2010), and the sequence capture targets were aligned to these clusters using blastn v2.2.31 (Camacho, et al. 2009). The sequences corresponding to each target were extracted for each sample and aligned using mafft v7.313 (Katoh and Standley 2013) and all 7,139 sequence alignments (1 per target) were concatenated. RAxML v8.2.11(Stamatakis 2014) was used to generate a maximum likelihood phylogeny using 100 rapid bootstrap replicates and the GTRCAT model of sequence evolution. Results suggested sufficient phylogenetic resolution, but one unexpected finding was the placement of *T. carnifex* as the sister lineage to *T. dobrogicus* (Supplementary Fig. S1a). Yet, in our main experiment, *T. carnifex* was more closely related to *T. macedonicus* (see Results). The fact that the *T. carnifex* sample used for transcriptome sequencing originated from close to the documented hybrid zone with *T. dobrogicus* (Arntzen, et al. 2014; Wielstra, Sillero, et al. 2014) suggests that substantial interspecific gene flow might underlie this relationship. To further explore this scenario we obtained transcriptomes from two additional *T. carnifex* individuals, sampled away from the hybrid zone with *T. dobrogicus*, representing the distinct Balkan and Italian mtDNA clades (Canestrelli, et al. 2012; Wielstra, et al. 2013). We processed these two individuals as above and reran RAxML, replacing the *T. carnifex* sample from the hybrid zone, and found that *T. carnifex* was recovered as the sister lineage to *T. macedonicus* (Supplementary Fig. S1b). Assuming that the *T. carnifex-T. macedonicus* relationship is correct, this phylogenetic shift reflects both the general risk of single-exemplar sampling (Spinks, et al. 2013) and the distorting influence that interspecific gene flow can have on phylogenetic inference (Leaché, et al. 2014). These findings support our decision to include multiple samples per species and to exclude samples from near known hybrid zones in our main experiment.

**Text S2: Processing of Sequence Capture Data**

*Reference assembly* – Sequence reads from the sample with the most reads (*T. carnifex* 292 with 3,937,346 read pairs) were used *de novo* to assemble target sequences for each target region. Trimmomatic v0.36 (Bolger, et al. 2014) was first used to remove adapter contamination and to trim leading bases with scores < 5, trailing bases with scores < 15, also employing a 4bp sliding window from 5’ to 3’, trimming the window and downstream sequence when the average quality of the window dropped < 20. Reads < 40bp were discarded. Trimmed reads were input into PEAR v0.9.10 (Zhang, et al. 2014) to merge overlapping paired end reads into longer single-end fragments with the following settings: p-value = 0.01, minimum assembly length = 50, statistical method = OES, using empirical frequencies = YES, quality score threshold = 0, minimum overlap = 10, and scaling method = scaled score.

Unmerged reads and merged read pairs were input into the assembly by reduced complexity (ARC) pipeline (Hunter, et al. 2015), which performs alternating tasks of mapping reads to target sequences, followed by per-target *de novo* assembly of mapped reads, replacing the original target sequences with the target assembly at each iteration. Six iterations were performed to generate a set of reference contigs assembled from reads relevant to each target region. A single assembled contig was then selected for each original target region by means of reciprocal best blast hit (RBBH) (Rivera, et al. 1998). These RBBHs were then blasted against one another to determine self-complementary regions, which can indicate chimeric assembly regions, and regions found to be similar to other target regions were trimmed to the nearest terminus of the contig (McCartney-Melstad, et al. 2016). This set of chimera-trimmed RBBHs was used as a target reference assembly for all downstream analyses.

*QC, SNP calling and genotyping* – Adapter contamination from library DNA inserts < 150bp was removed from reads using skewer v0.2.2 (Jiang, et al. 2014). Reads were mapped to the reference assembly using BWA-MEM v0.7.15-r1140 (Li 2013). Picard tools v2.9.2 (https://broadinstitute.github.io/picard/) was used to add read group information and mark PCR duplicates, and GATK v3.8 was used to generate gVCFs for each sample using HaplotypeCaller. GenotypeGVCFs was used for groups of samples (crested newts or crested + marbled newts, depending on the analysis) to call SNPs/genotypes, removing SNPs flagged by the following hard filters: QD < 2, MQ < 40, FS > 60, MQRankSum < -12.5, ReadPosRankSum < -8, QUAL < 30 (DePristo, et al. 2011; Poplin, et al. 2017).

The *de novo* assembly followed by RBBH approach is susceptible to the inclusion of paralogous loci as putatively single-copy targets. Because fixed differences between paralogues will appear as consistently heterozygous SNPs, we next attempted to remove paralogous targets from our dataset through the use of a Hardy Weinberg Equilibrium (HWE) filter for heterozygote excess. Heterozygote excess p-values were calculated for every SNP using vcftools 0.1.15 (Danecek, et al. 2011), and any target containing at least one SNP with a heterozygote excess p-value less than 0.05 was removed from downstream analysis.

*Reference assembly and genotyping* – A total of 4,932,636 reads (including 2,579,319 merged read pairs with an average length of 196bp) were used as input in the ARC assembly pipeline. After six iterations of mapping and assembly, 6,970 targets finished with an average of 295 reads apiece (median=167, sd=1,152), and 6,686 of the original targets had RBBHs to the assembly. After self-blast and trimming to remove potentially chimeric assemblies, a reference assembly of 5,593,497bp was generated for subsequent read mapping and SNP calling.

A median of 44.1% of trimmed reads aligned to the reference assembly (min=41.0%, max=50.5%), and an average of 22.6% of mapped reads were flagged as PCR duplicates, yielding a median unique reads on target of 34.2% (min=31.3%, max=39.4%). For the 23-sample dataset including the two marbled newt species, a total of 370,007 SNPs were recovered that passed hard filters. Of the 6,686 starting targets, 798 were found to contain at least 1 SNP with a HWE heterozygote excess p-value less than 0.05 and were removed. For the 21-sample dataset that did not contain the marbled newts, a total of 286,691 SNPs passed the hard filters and 814 targets were removed because they failed the HWE filter. Pairwise F84 divergences calculated with Phylip 3.697 (Felsenstein 1989) and based on the 23-sample dataset (including all *Triturus* species) are provided in Supplementary Table S2. The highest intraspecific divergence was observed between the Italian and Balkan clades comprising *T. carnifex*.

**Text S3: Phylogenetic Analyses**

*Data concatenation with RAxML* – RAxML version 8.2.11 (Stamatakis 2014) was used to infer phylogenies from concatenated alignments of SNPs. All biallelic SNPs in the 23-sample dataset that had genotype qualities of at least 20, that were present in at least 50% of the samples, and that fit RAxML’s definition of variable (133,601 SNPs total across 5,866 different targets) were used for maximum likelihood phylogenetic analysis. 100 rapid bootstrap replicates and 20 maximum likelihood searches were conducted with the ASC_GTRGAMMA model with Lewis ascertainment correction for SNP analysis (Lewis 2001). The resulting phylogeny with bootstrap support values was plotted in R using phytools (Revell 2011).

The mean depth of passing genotype calls across all samples was 42.4X, and median per-site missingness was 4.3%, which corresponds to one sample out of 23 missing data for a site (mean=10.1%, sd=14.0%). All crested newt species (for which three individuals were included) were recovered as monophyletic, and all bootstrap values on the tree were 100 (Fig. 3). The longest branch was between the marbled and crested newts and was used to root the tree. Within the crested newts, *T. ivanbureschi* was the sister lineage to a clade consisting of *T. anatolicus* and *T. karelinii*. The remaining four species were the sister-group to this assemblage, with *T. carnifex* most closely related to *T. macedonicus* and *T. cristatus* most closely related to *T. dobrogicus*. Since the monophyly of all species was strongly supported, species designations were fixed for subsequent species-tree inference.

*Gene-tree summary with ASTRAL* – ASTRAL v5.6.1 was used to estimate the crested newt phylogeny and to explore gene-tree discordance, presumably derived primarily from incomplete lineage sorting from a collection of gene-trees (Mirarab, et al. 2014; Sayyari and Mirarab 2016; Zhang, et al. 2017). No marbled newts were included because estimating terminal branch lengths is not possible for species with a single representative (note that the reciprocal monophyly of crested and marbled newts is well established (Arntzen, et al. 2007; Espregueira Themudo, et al. 2009; Wielstra, Arntzen, et al. 2014) and also strongly supported by our concatenated RAxML analysis). Separate polymorphic SNP alignments were first generated for each target using SnpSift 4.3 (Ruden, et al. 2012) and PGDSpider 2.1.1.2 (Lischer and Excoffier 2012), omitting SNPs with > 50% missing data across the 21 crested newt samples and removing targets that contained one or more samples with 100% missing data across the target using trimal v1.4.1 (Capella-Gutiérrez, et al. 2009). RAxML v8.2.11 (Stamatakis 2014) was used to infer a maximum likelihood gene-tree for each target with the ASC_GTRGAMMA model and Lewis ascertainment bias correction (Lewis 2001).

After setting genotypes with quality scores less than 20 to missing data and filtering out sites with > 50% missing data, a total of 143,571 SNPs remained across 5,861 targets to build gene-trees. After removing targets that contained samples with 100% missing data and removing sites that RAxML determined to be monomorphic, maximum likelihood gene-trees were built for 5,610 targets. These gene-trees were used as input into ASTRAL, constraining the seven crested newt species to be monophyletic (as supported by our concatenated RAxML analysis) and outputting local posterior probabilities and inferring terminal branch lengths. Midpoint rooting was used to determine the root. ASTRAL yielded a final normalized quartet score of 0.63. The same topology as in the concatenated RAxML analysis was recovered, with local posterior probabilities of 1 for all nodes (Fig. 4a). Branch lengths in ASTRAL are measured in coalescent units and indicate the degree of discordance among gene-trees (within taxa for terminal branches and among taxa for internal branches). The longest terminal branch was recovered for *T. macedonicus*, and the shortest belonged to *T. anatolicus*. The shortest internal branches were those separating the sister lineages *T. carnifex* + *T. macedonicus* from *T. cristatus* + *T. dobrogicus*.

*Species-tree estimation with SNAPP* – The coalescent species-tree inference method SNAPP v1.3.0 was used to infer the crested newt species-tree from biallelic SNPs (Bryant, et al. 2012). Marbled newts were not included because they introduce a long internal branch that can render parameter estimation inaccurate and splits between them and crested newts is not a primary goal of our paper. Polymorphic biallelic SNPs with genotype phred scores ≥ 20 across all 21 crested newts were first collected. Then, a single SNP from each of the 5,581 remaining loci was randomly selected to reduce the impacts of physical genetic linkage. These SNPs were used as input into SNAPP within the BEAST v2.4.8 environment (Bouckaert, et al. 2014) with the following parameters: species assignment=7 respective species, mutation rate U=1.0, mutation rate V=1.0, coalescence rate=10.0 (and sampled), use log likelihood correction=True, lambda prior=Gamma (initial=10[0.0,inf]) with alpha=2.0 and beta=200.0, snapprior.alpha=1.0, snapprior.beta=250.0, snapprior.kappa=1.0, snapprior.lambda=10.0 (and sampled), snapprior.rateprior=gamma, chain length=10,000,000, store every=1000 (and logging every 1000), and pre burnin=0. A 10% burnin was used and convergence and mixing were assessed with Tracer v1.7.1 (Rambaut, et al. 2018). ESS values for all parameters were > 400. A maximum clade credibility tree was constructed with common ancestor heights using TreeAnnotator v2.4.8 (Bouckaert, et al. 2014). Note that BEAST infers the root as part of the analysis. The same topology as in the RAxML and ASTRAL analyses was recovered (Fig. 4b). All posterior probabilities were 1, except for the node subtending *T. carnifex* + *T. macedonicus*, which was 0.99.

*Molecular dating with SNAPP* – A time-calibrated phylogeny was estimated with SNAPP using the same input SNP file as above. For calibration we interpreted the origin of the Adriatic Sea at the end of the Messinian Salinity Crisis at 5.33 million years ago (Krijgsman, et al. 1999) as the vicariance event causing the *T. carnifex* versus *T. macedonicus* split (Arntzen, et al. 2007; Wielstra and Arntzen 2011) and set the age of their most recent common ancestor to a uniform distribution between 5.32 and 5.34 million years ago (Stange, et al. 2018). Input XML files for divergence time estimation were prepared using snapp_prep.rb (https://github.com/mmatschiner/snapp_prep). We recognize that this is only a rough approximation given a single, biogeographically-informed date calibration point, and use it primarily to estimate the closeness in time of the crested newt radiation. The output tree from the original, undated SNAPP analysis was used as a starting tree, scaling the entire tree so that the starting age of the calibration node was 5.33 million years ago. The topology was fixed to that recovered by the original SNAPP analysis, and dates of remaining nodes were estimated using 1,000,000 MCMC steps, sampling every 500 steps and removing a 10% burn-in. ESS values for parameters were confirmed > 400 with Tracer. A maximum clade credibility tree with median node heights was generated with TreeAnnotator (Fig. 5).

**Text S4: Comparison with full mtDNA-based phylogeny**

MtDNA has proven misleading at both recent (Rodríguez, et al. 2017) and deeper (Veith, et al. 2018) nodes in the Salamandridae phylogeny and our genome-enabled phylogeny shows a highly supported deviation with a previous full mtDNA (i.e. single marker) phylogeny as well (Wielstra and Arntzen 2011). The deviation concerns the relationship among the three species constituting the ‘*T. karelinii*-group’; we here recover *T. anatolicus* as the sister lineage to *T. karelinii*, rather than to *T. ivanbureschi* as suggested by mtDNA (Supplementary Fig. S2). While such gene-tree discordance could reflect incomplete lineage sorting of mtDNA (Platt, et al. 2018), we consider ancient mtDNA introgression more likely, as *T. ivanbureschi* and *T. anatolicus* show geographically extensive introgressive hybridization today (Wielstra, et al. 2017). A scenario of ancient introgression is in line with the high degree of gene-tree/species-tree discordance in the nuclear genome in *T. anatolicus*, as suggested by the short branch in the ASTRAL tree (Fig. 4a). However, as all members of the ‘*T. karelinii*-group’ possess an identical number of trunk vertebrae, the mtDNA-nuDNA mismatch does not influence our interpretation of character evolution (Supplementary Fig. S3). The calibrated nuclear DNA-based (Fig. 5) and mtDNA-based phylogenies agree that cladogenesis among crested newts occurred over a relatively brief time window. However, mtDNA-based dates are older (cf. Table 2 in Wielstra and Arntzen 2011). This could simply reflect the differences in the dating method and the (slight) differences in the calibration scheme applied, but it is well-known that divergence times derived from individual gene-trees, and particularly from mtDNA, can be overestimates of lineage divergence (McCormack, et al. 2011).

**Text S5: Inference of changes in the number of trunk vertebrae**

The number of trunk vertebrae (NTV) in crested newts is characterized by a punctuated continuous character state distribution, with modal values for NTV in the range of 13-16 (for convenience an NTV count of 16 was used for *T. dobrogicus*, but note that NTV = 17 also occurs at roughly equal frequency in this species, which does not influence our interpretation). We consider NTV = 12, as observed in the sister lineage the marbled newts (the *T. marmoratus*-group), as well as the most closely related genus *Lissotriton*, to be the ancestral state (Arntzen, et al. 2015; Veith, et al. 2018). We applied the parsimony criterion to infer changes in NTV along all possible crested newt topologies (Supplementary Fig. S3). The program PAUP* (Swofford 1998) was used to allocate character state gains and losses over the tree, under ACCTRAN as well as DELTRAN optimization.

**Supplementary Figures S1-S3**

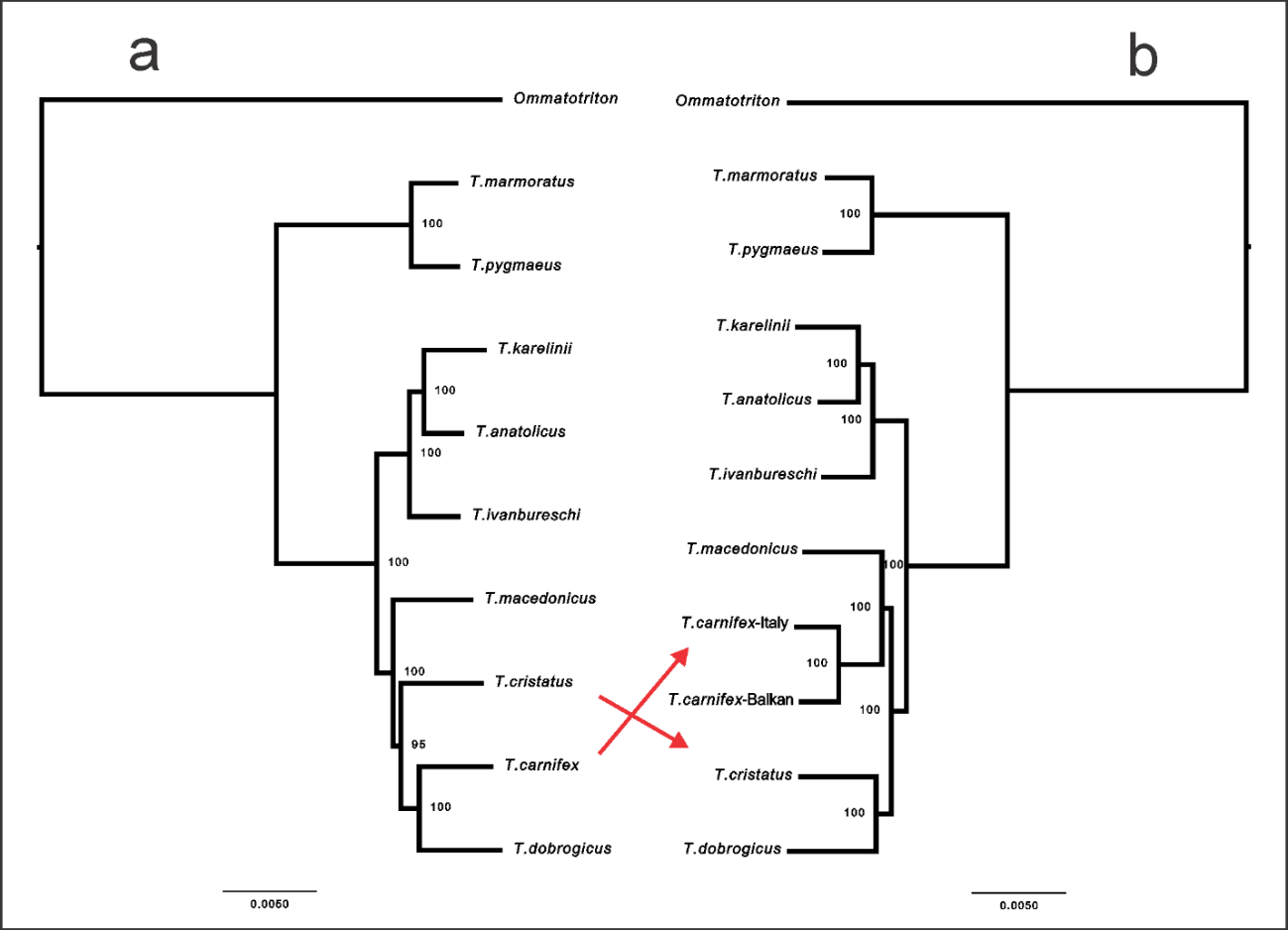

**Fig. S1. *Triturus* newt phylogenies based on based on data concatenation of transcriptome data with RAxML.** In a) a *T. carnifex* individual is included that is suspected to be admixed with *T. dobrogicus* and in b) this is replaced by two other *T. carnifex* individuals assumed to not be affected by genetic admixture, one from the Balkans and one from Italy, away from the contact zone with *T. dobrogicus*. Note the differences in sister species relationships (reflected by red arrows), with the phylogeny in b) being in full agreement with the one based on target capture data (Fig. 3; Fig. 4).

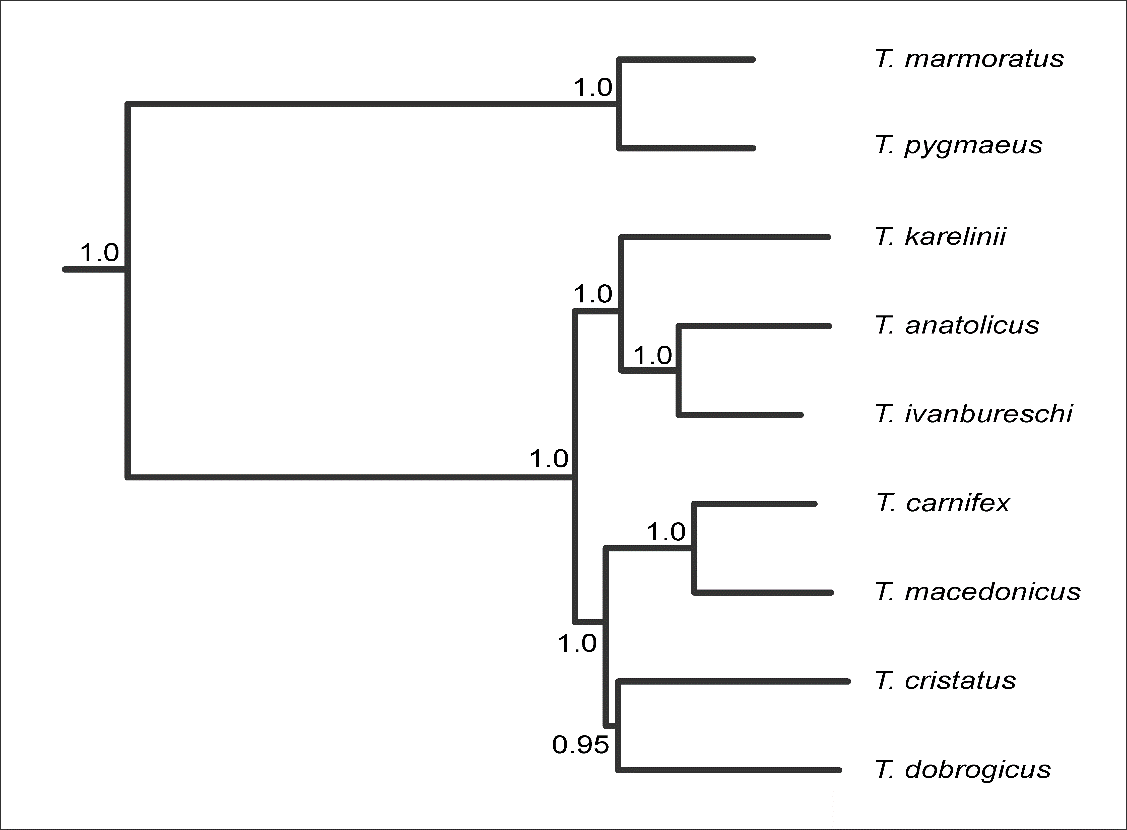

**Fig. S2. Full mtDNA phylogeny for *Triturus*.** The genome-enabled *Triturus* phylogeny (Fig. 3; Fig. 4) deviates from the phylogeny based on full mtDNA (taken from (Wielstra and Arntzen 2011)) for the species relationships in the *T. karelinii*-group of crested newts (with *T. anatolicus* being sister to *T. karelinii* rather than *T. ivanbureschi*). Numbers at nodes indicate posterior probabilities. Note the relatively low support for the sister relationship between *T. cristatus* and *T. dobrogicus*.

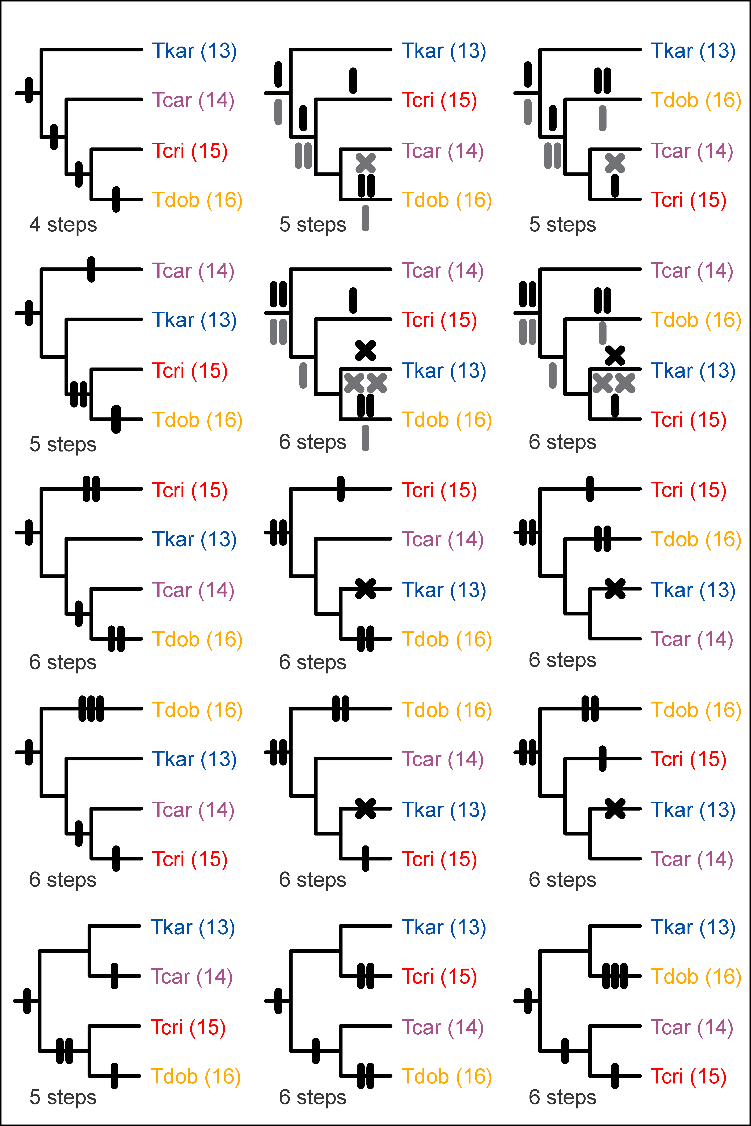

**Fig. S3. All 15 topologies possible for a fully bifurcating phylogeny of the four crested newt body builds.** Abbreviations: Tkar = *T. karelinii*-group; Tcar = *T. carnifex*-group; Tcri = *T. cristatus*; Tdob = *T. dobrogicus*. The number of trunk vertebrae (NTV) for each body build is provided in parentheses. A bar represents an NTV addition and a cross a deletion. NTV changes were inferred under the parsimony criterion, considering NTV = 12 as the ancestral character state for *Triturus* (see Supplementary Text S5). Results under ACCTRAN and DELTRAN optimization were identical for 11 topologies; for the four ones that deviated, character state changes under DELTRAN optimization are in black and above and under ACCTRAN optimization in grey and below branches. The top left topology corresponds to the *Triturus* species tree (Fig. 3; Fig. 4).

**Supplementary Tables S1-S2**

**Table S1. Sampling details.** Individuals are identified with a code that refers to complete specimens (ID starting with ZMA) or tail tips (remaining samples). All material is stored at Naturalis Biodiversity Center, Leiden, The Netherlands.

| ***Target capture*** |  |  |  |  |
| --- | --- | --- | --- | --- |
| **ID** | **Species** | **Locality** | **Latitude** | **Longitude** |
| 5017 | *Triturus marmoratus* | France: Jublains | 48.252 | -0.473 |
| 5016 | *Triturus pygmaeus* | Portugal: Serra de Monchique | 37.335 | -8.506 |
| 4729 | *Triturus ivanbureschi* | Bulgaria: Ostar Kamak | 41.878 | 25.853 |
| 1814 | *Triturus ivanbureschi* | Turkey: Karakadılar | 40.010 | 26.940 |
| 1788 | *Triturus ivanbureschi* | Turkey: Bozdağ | 38.367 | 28.103 |
| 1847 | *Triturus anatolicus* | Turkey: Abanta Gölu | 40.612 | 31.288 |
| 1889 | *Triturus anatolicus* | Turkey: Gölköy | 40.083 | 33.347 |
| 1985 | *Triturus anatolicus* | Turkey: Çakırlı | 40.446 | 37.483 |
| 2105 | *Triturus karelinii* | Ukraine: Nikita | 44.538 | 34.243 |
| 6719 | *Triturus karelinii* | Azerbaijan: Altiagac | 40.854 | 48.935 |
| RMNH RenA 46931-2390 | *Triturus karelinii* | Iran: Qu’Am Shahr | 36.436 | 52.803 |
| ZMA9108-405 | *Triturus carnifex* | Italy: Fuscaldo | 39.417 | 16.033 |
| ZMA9145-292 | *Triturus carnifex* | Italy: Pisa | 43.717 | 10.400 |
| ZMA9132-312 | *Triturus carnifex* | Slovenia: Kramplje | 45.733 | 14.500 |
| 3247 | *Triturus macedonicus* | Montenegro: Bjeloši | 42.374 | 18.907 |
| 3275 | *Triturus macedonicus* | Albania: Bejar | 40.429 | 19.850 |
| 3775 | *Triturus macedonicus* | Greece: Kerameia | 39.562 | 22.081 |
| 4485 | *Triturus cristatus* | Bulgaria: Montana | 43.416 | 23.222 |
| 3686 | *Triturus cristatus* | Romania: Budeni | 45.768 | 26.839 |
| ZMA9167-355 | *Triturus cristatus* | Romania: Virfuri | 46.283 | 22.467 |
| ZMA9083-512 | *Triturus dobrogicus* | Hungary: Alap | 46.800 | 18.683 |
| ZMA9172-720 | *Triturus dobrogicus* | Croatia: Zupanja | 45.083 | 18.700 |
| 2377 | *Triturus dobrogicus* | Romania: Mǎcin | 45.251 | 28.121 |
| ***Transcriptomes*** |  |  |  |  |
| **ID** | **Species** | **Locality** | **Latitude** | **Longitude** |
| 6720 | *Triturus marmoratus* | Portugal: Valongo | 41.168 | -8.500 |
| 6721 | *Triturus pygmaeus* | Portugal: Serra de Monchique | 37.335 | -8.506 |
| 6722 | *Triturus karelinii* | Azerbaijan: Katex | 41.646 | 46.543 |
| 6723 | *Triturus anatolicus* | Turkey: Hacılar | 41.495 | 32.088 |
| 6724 | *Triturus ivanbureschi* | Turkey: Keşan | 40.924 | 26.635 |
| 6725 | *Triturus carnifex* | Croatia: Prkovac | 45.569 | 16.094 |
| 6726 | *Triturus carnifex* | Croatia: Radetići | 45.146 | 13.842 |
| 6727 | *Triturus carnifex* | Italy: Viterbo | 42.703 | 13.325 |
| 6728 | *Triturus macedonicus* | Montenegro: Ceklin | 42.367 | 18.982 |
| 6729 | *Triturus cristatus* | France: Belgeard | 48.259 | -0.574 |
| 6730 | *Triturus dobrogicus* | Serbia: Sremski Karlovski | 45.175 | 19.991 |
| 6731 | *Ommatotriton nesterovi* | Turkey: Hürriyet | 40.276 | 28.650 |

**Table S2. Inter- and intraspecific divergence in *Triturus* newts.** Shown are pairwise F84 divergences calculated with Phylip. Intraspecific distances are in italics. IDs correspond to Supplementary Table S1.

| *T. dobrogicus* | | | *T. cristatus* | | | *T. macedonicus* | | | *T. carnifex* | | | *T. karelinii* | | | *T. anatolicus* | | | *T. ivanbureschi* | | | *T. pygmaeus* | *T. marmoratus* |  |
| --- | --- | --- | --- | --- | --- | --- | --- | --- | --- | --- | --- | --- | --- | --- | --- | --- | --- | --- | --- | --- | --- | --- | --- |
| **2377** | **720** | **512** | **355** | **3686** | **4485** | **3775** | **3275** | **3247** | **312** | **292** | **405** | **2390** | **6719** | **2105** | **1985** | **1889** | **1847** | **1788** | **1814** | **4729** | **5016** | **5017** |  |
| 0.70 | 0.69 | 0.71 | 0.69 | 0.72 | 0.71 | 0.75 | 0.72 | 0.74 | 0.71 | 0.75 | 0.74 | 0.73 | 0.74 | 0.71 | 0.72 | 0.72 | 0.68 | 0.72 | 0.72 | 0.72 | 0.10 | - | **5017** |
| 0.68 | 0.67 | 0.69 | 0.67 | 0.70 | 0.69 | 0.73 | 0.70 | 0.72 | 0.69 | 0.73 | 0.72 | 0.71 | 0.72 | 0.69 | 0.70 | 0.70 | 0.66 | 0.71 | 0.70 | 0.70 | - |  | **5016** |
| 0.22 | 0.21 | 0.23 | 0.21 | 0.22 | 0.21 | 0.24 | 0.23 | 0.24 | 0.23 | 0.25 | 0.25 | 0.14 | 0.14 | 0.14 | 0.11 | 0.11 | 0.08 | *0.03* | *0.02* | - |  |  | **4729** |
| 0.22 | 0.21 | 0.23 | 0.21 | 0.22 | 0.21 | 0.24 | 0.23 | 0.24 | 0.23 | 0.25 | 0.25 | 0.14 | 0.14 | 0.14 | 0.11 | 0.11 | 0.08 | *0.02* | - |  |  |  | **1814** |
| 0.22 | 0.22 | 0.24 | 0.21 | 0.22 | 0.22 | 0.25 | 0.23 | 0.25 | 0.24 | 0.26 | 0.26 | 0.15 | 0.15 | 0.14 | 0.12 | 0.12 | 0.09 | - |  |  |  |  | **1788** |
| 0.20 | 0.20 | 0.21 | 0.19 | 0.20 | 0.19 | 0.22 | 0.20 | 0.22 | 0.21 | 0.23 | 0.23 | 0.10 | 0.10 | 0.09 | *0.03* | *0.02* | - |  |  |  |  |  | **1847** |
| 0.22 | 0.22 | 0.24 | 0.21 | 0.22 | 0.22 | 0.25 | 0.23 | 0.25 | 0.24 | 0.26 | 0.26 | 0.12 | 0.12 | 0.11 | *0.03* | - |  |  |  |  |  |  | **1889** |
| 0.22 | 0.22 | 0.23 | 0.21 | 0.22 | 0.21 | 0.25 | 0.23 | 0.25 | 0.23 | 0.25 | 0.25 | 0.11 | 0.11 | 0.11 | - |  |  |  |  |  |  |  | **1985** |
| 0.23 | 0.23 | 0.25 | 0.22 | 0.23 | 0.23 | 0.26 | 0.24 | 0.25 | 0.25 | 0.26 | 0.27 | *0.01* | *0.01* | - |  |  |  |  |  |  |  |  | **2105** |
| 0.24 | 0.23 | 0.25 | 0.23 | 0.24 | 0.23 | 0.26 | 0.24 | 0.26 | 0.25 | 0.27 | 0.27 | *0.02* | - |  |  |  |  |  |  |  |  |  | **6719** |
| 0.24 | 0.23 | 0.25 | 0.23 | 0.24 | 0.23 | 0.26 | 0.24 | 0.26 | 0.25 | 0.27 | 0.27 | - |  |  |  |  |  |  |  |  |  |  | **2390** |
| 0.19 | 0.19 | 0.21 | 0.21 | 0.21 | 0.21 | 0.22 | 0.20 | 0.22 | *0.08* | *0.03* | - |  |  |  |  |  |  |  |  |  |  |  | **405** |
| 0.19 | 0.18 | 0.20 | 0.21 | 0.21 | 0.21 | 0.22 | 0.20 | 0.21 | *0.07* | - |  |  |  |  |  |  |  |  |  |  |  |  | **292** |
| 0.17 | 0.17 | 0.18 | 0.19 | 0.19 | 0.19 | 0.20 | 0.18 | 0.19 | - |  |  |  |  |  |  |  |  |  |  |  |  |  | **312** |
| 0.19 | 0.19 | 0.21 | 0.19 | 0.20 | 0.20 | *0.06* | *0.04* | - |  |  |  |  |  |  |  |  |  |  |  |  |  |  | **3247** |
| 0.18 | 0.17 | 0.19 | 0.17 | 0.18 | 0.18 | *0.04* | - |  |  |  |  |  |  |  |  |  |  |  |  |  |  |  | **3275** |
| 0.19 | 0.19 | 0.21 | 0.19 | 0.20 | 0.20 | - |  |  |  |  |  |  |  |  |  |  |  |  |  |  |  |  | **3775** |
| 0.15 | 0.16 | 0.17 | *0.04* | *0.05* | - |  |  |  |  |  |  |  |  |  |  |  |  |  |  |  |  |  | **4485** |
| 0.16 | 0.16 | 0.17 | *0.05* | - |  |  |  |  |  |  |  |  |  |  |  |  |  |  |  |  |  |  | **3686** |
| 0.15 | 0.16 | 0.17 | - |  |  |  |  |  |  |  |  |  |  |  |  |  |  |  |  |  |  |  | **355** |
| *0.02* | *0.03* | - |  |  |  |  |  |  |  |  |  |  |  |  |  |  |  |  |  |  |  |  | **512** |
| *0.03* | - |  |  |  |  |  |  |  |  |  |  |  |  |  |  |  |  |  |  |  |  |  | **720** |
| - |  |  |  |  |  |  |  |  |  |  |  |  |  |  |  |  |  |  |  |  |  |  | **2377** |
